# Sex-Dependent Effects of Chronic Stress During Adolescence on Cognitive Bias and Functional Connectome in Young Adult Rats

**DOI:** 10.64898/2026.03.18.712614

**Authors:** Twain Dai, Liz Jaeschke-Angi, Marissa Penrose-Menz, Tim Rosenow, Jennifer Rodger

**Author notes:** **Correspondence:** Twain Dai.

## Abstract

Negative cognitive biases in depression are more pronounced in females than in males. This sex difference emerges during adolescence, a sensitive developmental stage when chronic stress exposure increases the risk of depression in adulthood. Neuroimaging studies in adult depression suggest that sex differences in negative bias are associated with aberrant functional connectivity. However, the neurobiology linking adolescent stress to sex-specific cognitive bias and resting-state network reorganization in adults remains poorly understood. The study aimed to investigate the longitudinal effects of chronic restraint stress (CRS) during adolescence on cognitive bias and functional connectome in emerging adulthood. 28 Wistar rats (sex-balanced; aged five weeks on arrival) were trained on a judgment bias task with distinct tactile cues signaling differential rewards. Bias was quantified from responses to ambiguous probe trials. Following training, animals were randomly and equally assigned to CRS or control groups (sex-balanced). Offline resting-state functional MRI scans were conducted at adolescent baseline (pre-CRS) and again in adulthood (post-CRS), followed by probe trials to assess neural and behavioral changes. Behaviors were analyzed using linear mixed-effects models, and network connectivity was assessed using Network-Based R-Statistics. Both behavioral and network analyses identified sex-dependent differences in change from baseline to post-CRS. Specifically, females showed a greater tendency toward negative bias than males. Network analysis suggested female-specific reorganization within a subnetwork, spanning brainstem, limbic, striatal, insula, parahippocampal, orbitofrontal, and retrosplenial regions. These results highlight the translational importance of considering sex and distributed network organization when modelling how adolescent stress may shape later vulnerability to depression.

## Introduction

Adolescence is a critical developmental period during which the brain undergoes profound structural and functional refinements, particularly cortical-subcortical circuits that govern stress regulation and cognitive processing (1, 2). Recent large-scale connectomics studies reveal that different neural circuits undergo heterochronic refinement during the transition to adulthood (3, 4). While primary sensory-motor circuits reach functional stability in early childhood, higher-order affective and cognitive networks experience ongoing refinement throughout adolescence. This developmental mismatch highlights adolescence as a pivotal turning point where functional organization of the brain is uniquely sensitive to environmental adversities (5).

Chronic stress during adolescence is known to derail these developmental trajectories by disrupting the normative refinement of cortical-subcortical circuits, significantly increasing the risk of depression in adulthood (6). The resulting psychiatric vulnerability is behaviorally characterized by negative cognitive biases, a systematic tendency to prioritize the processing of aversive information over neutral or positive stimuli (7). These biases are not merely symptoms of affective dysfunction but are considered core factors that predict the onset and chronicity of depression (8, 9). This predictive relationship is notably stronger in females. Longitudinal behavioral evidence demonstrates that the interaction between negative cognitive biases and stress more robustly affects depression trajectories in females than in males during adolescence (10).

The sex difference in negative cognitive biases is mirrored by distinct neural signatures. Specifically, adolescent females with depression exhibit unique recruitment of the posterior cingulate and supramarginal gyrus when faced with negative distractors, a response absent in their male counterparts (11). Furthermore, the neural correlates of cognitive bias are divergent during memory processing: while adolescent females with depression exhibit amygdala hyperactivity during negative recall, male counterparts are characterized by reduced hippocampal engagement during positive recall (12). Although human neuroimaging studies in adult depression suggest that sex differences in negative cognitive bias are associated with aberrant functional connectivity, the neurobiology linking adolescent chronic stress to sex-dependent cognitive bias and resting-state network reorganization remains poorly understood. Gaining a deeper understanding of these mechanisms could inform novel, targeted strategies for preventing adult depression.

Although animal models cannot fully capture the heterogeneity and social complexity of adolescent adversity in humans, they offer unique advantages for examining the neurobiology of chronic stress under highly controlled conditions (13, 14). In rodents, chronic stress during adolescence has been shown to induce detrimental effects on affective behaviors in adulthood (15, 16). Similar to humans, a key manifestation is a shift toward negative bias, which can be objectively quantified using translational paradigms such as the judgment bias task (JBT). By measuring how animals interpret ambiguous cues to assess levels of optimism and pessimism, the JBT has demonstrated that adolescent stress induces a negative bias, mirroring that in human depression (17, 18). However, most preclinical research continues to prioritize males, leaving the neural mechanisms driving female-specific cognitive biases following chronic stress largely unexplored (19, 20). This knowledge gap is further magnified by a lack of longitudinal data mapping the sex-specific developmental trajectory of neural network as it unfolds from adolescent stress exposure through to adulthood (15).

Therefore, this study employed a longitudinal design to examine cognitive bias and whole-brain resting-state functional architecture within the same cohort of males and females, before and after adolescent chronic stress. The study aimed to determine whether this stress exposure was associated with sex-dependent alterations in cognitive bias and resting-state network organization in emerging adulthood.

## Methods

### Ethics Statement

Ethical approvals for the animal experiment (2023/ET000552) were obtained from the animal ethics committee of the University of Western Australia (UWA).

### Animals and Study Design

28 Wistar rats (14 females/14 males; aged 5 weeks and weighing 110 ∼ 200 g on arrival) were sourced from the Ozgene Animal Resources Centre, Western Australia. All rats were housed under a reverse 12-hour light-dark cycle with ad libitum food and water, in a temperature-controlled room located at UWA’s Animal Care Unit (Nedlands, WA). Rats aged 55 ∼ 70 days were categorized as adolescents, whereas those aged 70 ∼ 150 days were considered emerging adults (21). While the oestrous cycle was not explicitly tracked, the longitudinal nature of behavioral task and the focus on resting-state connectome reorganization likely mitigated the influence of acute hormonal fluctuations on primary outcomes. All animals (housed in pairs) were acclimatized to the new environment and handled for nine days after their arrival and prior to experiments.

Animals were randomly and equally assigned to two sex-balanced groups: control (7 females/7 males) and CRS groups (7 females/7 males). On Day 9 after their arrival, animals began single-housed and mild food restrictions overnight, with food pellets removed at 4 pm. From Day 10 onward, animals were trained on the JBT (*Figure 1*). Animals remained under mild food restrictions, with two hours of free access to food each day until pre-probe was completed.

**Figure 1.**
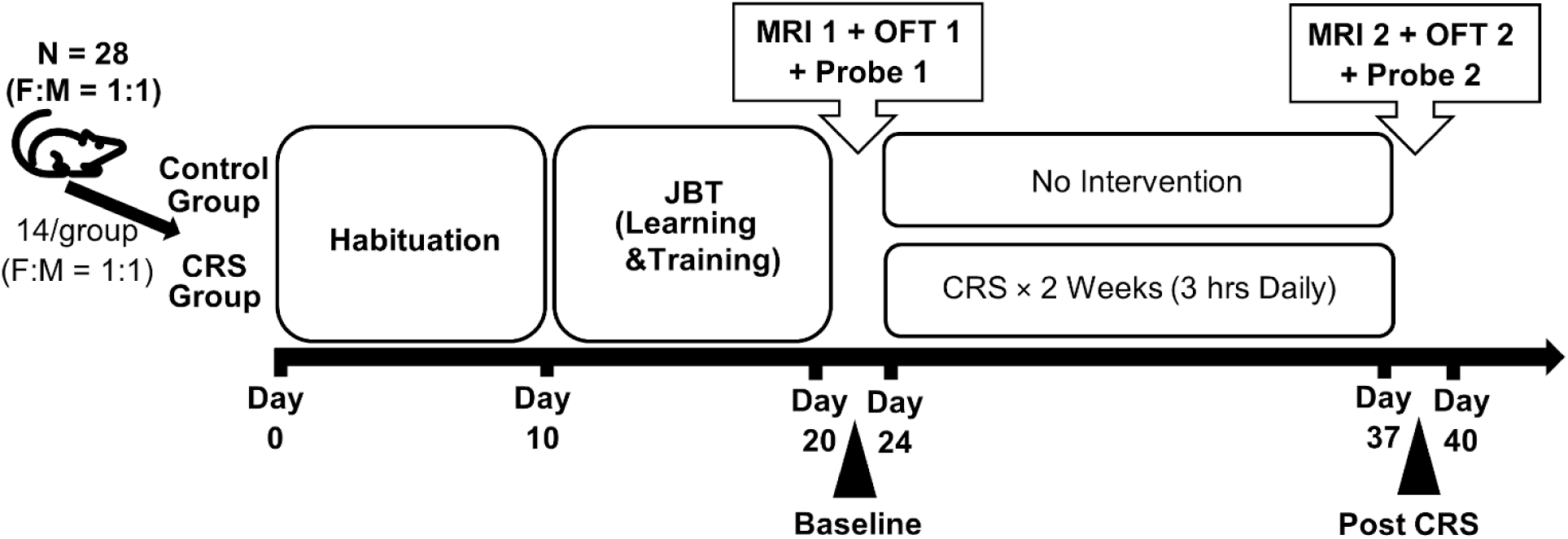
Study Design and Timeline. All animals were habituated to the new environment for nine days after their arrival and prior to experiments. All rats were trained on JBT, for up to nine days. Behavioral tests and MRI scan were conducted before and after 14 consecutive days of CRS. Investigators were not blinded to group allocation during CRS administration or behavioral testing, because knowledge of group assignment was required to deliver the appropriate procedures.

Animals were excluded from further experiments if they did not successfully complete JBT training within nine days. Once an animal was verified to have learned the task through the pre-probe test, it underwent a pre-OFT, which habituates rats to the testing arena. Following the pre-OFT, animals received no intervention until Day 21. Baseline MRI scan, OFT and JBT probe for all animals were conducted on Day 21, 22 and 23, respectively. From Day 24 onwards, animals in the CRS group underwent CRS for 14 consecutive days. The second MRI scan, OFT and JBT probe for all rats were carried out on Day 38, 39, and 40, respectively. For the baseline and post-CRS probes, food was removed overnight starting at 4 pm on Day 22 and 39, respectively.

### Judgment Bias Task

#### Task Design

The JBT in the present study was Go/Go task and adapted from Brydges and Hall (22). The task required animal’s responses to different tactile stimuli, olfactory and spatial cues, which were associated with different reward values. The task involved two reward values (large and small), two distinct tactile stimuli (coarse and fine), two distinct olfactory cues (cinnamon and coriander) and two distinct spatial cues (left and right), resulting in eight task combinations *(Supplementary Table 1)*. The assignment of task combinations was randomized equally across animals in each sex group and remained consistent for each animal throughout the experiment. The olfactory and location cues were employed to accelerate memory formation in learning the association between tactile stimuli and reward values.

#### Apparatus

The JBT apparatus (*Figure 2A*) was custom-built from acrylic in tint gray (The Plastic People, Osborne Park, WA). It consisted of three compartments: one start compartment (50 cm length × 50 cm width × 60 cm height) and two reward compartments (30 cm length × 15 cm width × 60 cm height). One removable barrier separated the start compartment from the reward compartments. A sandpaper (tactile stimuli) was attached to the center of the start compartment using Velcro tapes when required. One ceramic foraging bowl containing cheerios (rewards) was placed at the end of each reward compartment. One small muslin bag containing coriander or cinnamon was hung on each sidewall of each reward compartment.

**Figure 2.**
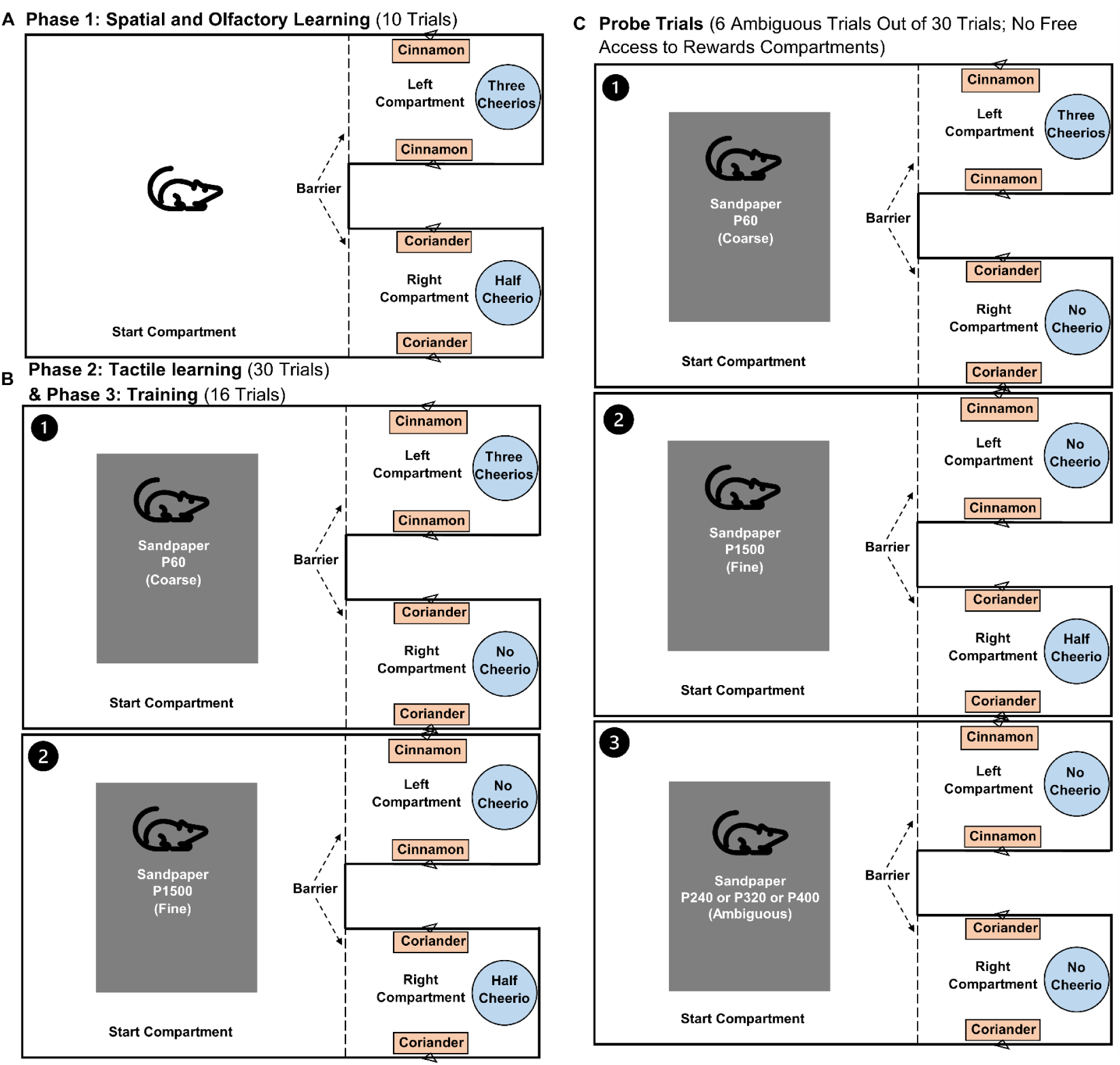
Learning, Training and Probe Protocols. The graph shows the procedure from a rat assigned task combination 1 (*Supplementary Table 1*). **A).** In the spatial and olfactory learning phase, the rat was trained to learn that high reward (three cheerios) was associated with cinnamon scent and left compartment, and low reward (half cheerio) was associated with coriander scent and right compartment; **B).** In the tactile learning phase, the same rat was further trained to learn that fine sandpaper indicated low reward and coarse sandpaper indicated high reward. In the training phase, an acrylic barrier was lowered to block movement between compartments as soon as the rat had made its choice; **C).** In the probe trials, one in every five trials was unrewarded. In these six unrewarded trials, three intermediate and ambiguous grades of sandpaper (P240, P320, P400) were randomly assigned and each presented twice as probes.

#### Procedures and Bias Measures

The JBT consisted of four phases: spatial and olfactory learning, tactile learning, training, and probe phases (*Figure 2*).

During the spatial and olfactory learning phase, animals were trained to learn the assigned association among reward values, spatial and olfactory cues. Different amounts of cheerios (rewards) were placed in two reward compartments. Animals were free to retrieve rewards from both compartments. This phase was considered complete when animals retrieved rewards from each compartment 5 times. Animals then proceeded to tactile learning the following day.

During the tactile learning phase, animals were further trained to learn the assigned association between reward values and tactile stimuli, in addition to the spatial and olfactory cues. Animals had free access to both reward compartments for a total of 30 trials. In each trial, only one compartment was baited, and a coarse (P60) or fine (P1500) sandpaper was secured to the floor in the center of the start compartment. The high and low reward trials were randomly assigned, with the rule that no more than two trials in a row were assigned the same size of reward. When animals completed 90% (27/30) of the trials within an hour, they proceeded to the training phase the following day.

The training phase was identical to the tactile learning phase, except that animals were only allowed to enter one reward compartment of their choice and prevented from entering the other reward compartment by placing an acrylic barrier following their choice. In addition, in one in every five trials, no reward was initially placed in either reward compartment; but animals were awarded a corresponding size of reward immediately after making the right choice, while no reward was given for an incorrect choice. This controlled any olfactory cues from cheerios that might guide decision-making and also served as reinforcement to enhance learning and reduce extinction. The training phase consisted of up to three training sessions in a day, with each training session containing 16 trials. Training continued until animals achieved 75% (12/16) accuracy in a training session.

All male rats (N = 14) completed the learning and training phases within four days, except for one that spent five days. Conversely, female rats (N = 12) took longer, with durations ranging from four to nine days. Two female rats did not complete the learning and training phases within nine days and were excluded from further experiments (Final N = 26; 12 females/14 males). Following the completion of training, animals underwent one pre-probe, one baseline probe, and one post-CRS probe session. The pre-probe was conducted the day after training was completed to verify that animals had learned assigned task combination. Baseline and post-CRS probes were carried out on Day 23 and 40, respectively. The probe phase was similar to the training phase, except that there were a total of 30 trials and one in every five trials was unrewarded. In these six unrewarded trials, three intermediate and ambiguous grades of sandpaper (P240, P320, P400) were randomly assigned and each presented twice as probes. The random presentation of three ambiguous tactile stimuli aimed to ensure that the animals did not learn that there were no rewards in ambiguous trials and always chose a reward compartment. All 26 animals completed 30 trials in the pre-probe, baseline and post-CRS probe sessions. Additionally, no refresher training was conducted during the CRS period.

Bias was assessed using response data from six ambiguous trials in the probe session that met the completion criteria, on an integer scale ranging from -3 to 3. A score of 3 represented an extremely positive bias, characterized by six high-reward responses; a score of 0 suggested a neutral bias, characterized by equal high-reward and low reward responses; whereas a score of -3 indicated an extremely negative bias, characterized by six low-reward responses. The overall response accuracy across 24 trials presenting distinct stimuli and the response accuracy associated with high and low reward outcomes were calculated. Additionally, the total time taken to complete a probe session (probe duration) was recorded.

### Chronic Restraint Stress

From Day 24 to Day 37, CRS was performed on a bench situated in one corner of a spacious animal holding room, with rats in the intervention group (6 females/7 males) placed in a transparent plastic tube facing the wall to mitigate visual distractions (23). Each session started between 12:00 and 13:00 PM, lasting for three hours daily to reduce the influence of circadian rhythm. Following each session, animals were immediately returned to their home cages. The control group (6 females/7 males) stayed in their home cages without undergoing CRS.

### Open Field Test and Behavioral Automation

OFT (24) was conducted in an open arena (50 cm length × 50 cm width × 60 cm height) to evaluate anxiety-like behaviors, with each animal tested individually. Animals were positioned initially in the center zone and then allowed to freely explore the arena for ten minutes. The apparatus was wiped with 70% ethanol to remove olfactory cues after each test session. Each session was recorded using a Samsung A70 mobile phone and saved in mp4 format for further processing and analysis. A total of 52 recordings were acquired at baseline and post-CRS. All original recordings were edited to be ten minutes in duration to remove irrelevant (non-test related) footage. The edited videos were then imported into DeepLabCut (25) to track animal movement by labeling seven body points (*Figure 3A*) and four corners of the testing arena (*Figure 3B*). Main steps involved in the behavioral labelling were: 1). Manually label 11 tracking points of 300 randomly selected video frames (20 frames × 15 videos); 2). Train a deep neural network using labelled frames; 3). Extract the coordinates of all tracking based on the trained network. For each recording, DeepLabCut automatically generated one individual CSV file containing the coordinates of 11 tracking points for all video frames, along with labelling confidence (ranging from 0 ∼ 100%) for each coordinate.

**Figure 3.**
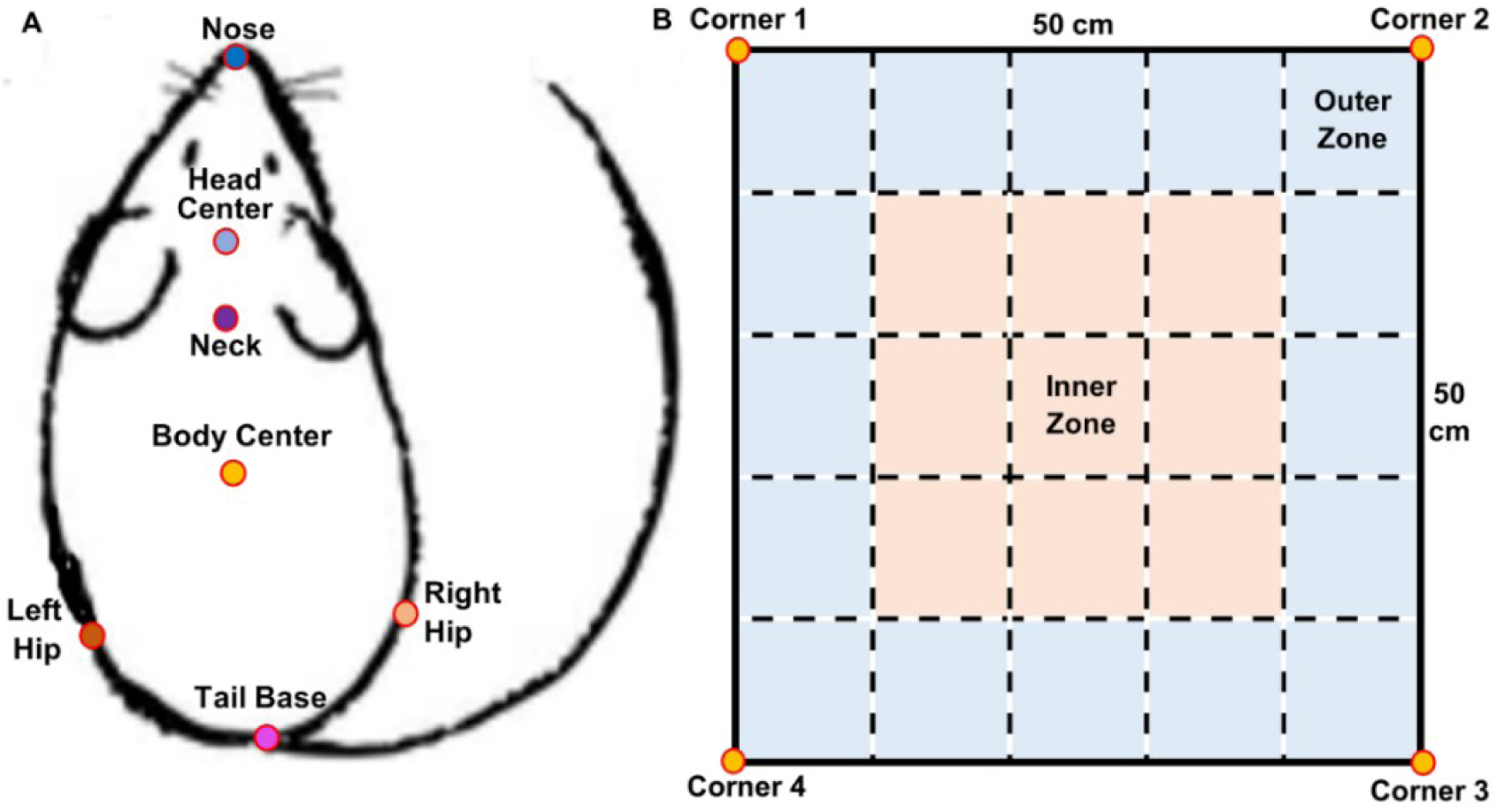
Labels Used to Train the Deep Neural Network. **A).** Rat with 7 body points; **B).** OFT outline with 4 corners. Beige denotes inner zone. Blue denotes the outer zone.

All DeepLabCut coordinate files derived from OFT recordings were imported into R 4.4.0. and processed with DLCAnalyzer (26) and in-house R scripts (*Supplementary Scripts*). For each video, the arena outline was first determined by the median coordinates of each corner point using DLCAnalyzer. Also, rat body points with a likelihood below 95% or located outside the testing arena were replaced through data interpolation. Then, the whole arena was virtually divided into a series of 10 x 10 cm blocks based on the arena outline using the in-house script. The outer zone consisted of 16 blocks (in blue, *Figure 3B*), while the inner zone contained 9 blocks (in beige, *Figure 3B*). The percentage of time spent in the outer zone during the first five minutes, second five minutes, and the total ten minutes was computed as a classical measure of anxiety-like behavior.

### MRI Acquisition and Batch Processing

Rats were scanned using Bruker Biospec 94/30 US/R pre-clinical MRI machine (Bruker BioSpin GmbH, Germany) operating at 9.4 T (400MHz; H-1). Animals were anesthetized with isoflurane and medetomidine during each MRI session; the anesthesia protocol was described and discussed thoroughly in previous work (23, 27). *Supplementary Table 2* summaries the parameters of two scanning sequences: T2w sMRI and rs-fMRI. Both raw images for each scan session were compiled in one ParaVision 6.3.4 package in the format of PvDatasets. A total of 23 animals completed both baseline and post-CRS scans. One control and one CRS female rat did not complete their baseline scans due to irregular respiratory rate, and one control male’s baseline package was corrupted due to technical issues. The batch processing of MRI data followed the same workflow as previous studies (27, 28). The workflow mainly included data organization to the Brain Imaging Data Structure (29), bias field correction, skull stripping, denoising, atlas registration to the Waxholm Space atlas (RRID:SCR_017124; 30), and functional connectivity computation. To compute the functional connectome of the whole brain for each animal, an in-house functional parcellation was mapped onto processed functional images (N = 46) to derive subject-specific time series for 13 functional networks (28). Fisher’s r-to-Z transformation (31) and partial correlation with Tikhonov regularization (default penalty strength of 0.1; 32, 33) was applied to these time series, resulting in 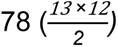 network connectivity/edges.

### Statistical Analysis

#### Behavioral Measures

Paired odds ratio (OR) for transition to negative bias from baseline to post-CRS was defined as the ratio of discordant transitions within each sex, using McNemar’s matched-pairs framework (34, 35). 95% confidence interval (CI) was computed on the log scale using the standard asymptotic variance for discordant pairs and exponentiated to return to the odds ratio scale. The sex difference in bias shifts was summarized as the ratio of paired odds ratios (ROR), with a log-scale CI calculated by combining the log-scale variances from the sex-specific paired ORs (35).

All behavioral measures were analyzed using Linear Mixed-Effects Models (LMMs) using the lme4 package (36) in R. These measures included bias score, response accuracy, probe duration, and the dwell time in the periphery of the OFT. To account for the longitudinal, repeated-measures structure of the data and the dependency within individual animals (N = 26), models were fitted with the formula: outcome ∼ sex × group × timepoint + (1 | animal). This included sex (male vs female), group (CRS vs control), and timepoint (baseline vs post-CRS) as fixed factors and their three-way interaction, with a random intercept for animal to control for repeated-measures variance. Model assumptions, including residual normality and homoscedasticity, were rigorously assessed using the performance package (37). For dependent variables violating these assumptions, a bootstrap method was applied to refit their models (36, 38). Significant three-way interactions were decomposed by assessing simple two-way interactions at each sex stratum, followed by simple main effects of timepoint, stratified by sex and group. Post-hoc comparisons were conducted using the rstatix (39) R packages, with Bonferroni corrections applied for multiple comparisons.

All measurements were presented as mean ± SD, unless otherwise specified. Effect sizes were reported with partial eta squared (η²p) and partial omega squared (ω²p). Since ω²p is widely viewed as lesser biased to η²p, effect sizes were evaluated based on ω²p and classified into four categories: < 0.02 as very small, 0.02 ∼ 0.13 as small, 0.13 ∼ 0.26 as medium, and ≥ 0.26 as large (40). Statistical significance was set as p < 0.05.

#### Network Connectivity

The network connectivity of the whole brain (78 edges) was analyzed using Network-Based R-Statistics with mixed-effects models (NBR; 41) to account for the longitudinal, repeated-measures structure of the data from 23 animals. NBR extends the original Network-Based Statistics framework (42) by incorporating linear mixed-effects models and applying permutation testing for family-wise error (FWE) control. This approach enables inference at the subnetwork level rather than relying solely on edgewise inference. First, a linear mixed-effect model was fitted to each edge using the same model structure as for the behavioral analysis in the previous section. Edges with p values below the initial significance threshold (p < 0.05) were then clustered into a subnetwork based on shared nodes. The term “subnetwork” refers specifically to the connected set of edges identified by NBR and not to a subset of the 13 predefined functional networks. Finally, the statistical significance of the subnetwork was evaluated against an empirical null distribution of subnetwork size and strength generated through 5000 permutations. P values were corrected for FWE and calculated as the proportion of permutations in which the maximal component statistic equaled or exceeded the observed values. The identified subnetwork was visualized as a chord diagram, and the corresponding null distributions of subnetwork size and strength were shown as density plots. Significant three-way interactions were further decomposed by assessing simple two-way interactions at each sex stratum, followed by simple main effects of timepoint within each sex-by-group stratum. These follow-up analyses followed the same procedure described above, but were restricted to the identified subnetwork.

To explore whether the identified subnetwork could be represented as a scalar outcome, the mean connectivity across edges within the subnetwork was computed and analyzed using a follow-up linear mixed-effects model. This scalar summary did not reproduce the significant three-way interaction, suggesting that a simple average of edge values did not adequately capture the subnetwork pattern identified by NBR.

Scripts and workspace for statistical analysis can be found in *Supplementary Script* and *Workspace*.

## Results

### Bias Measures

Following CRS, female rats showed higher odds of transitioning towards a negative bias (paired OR = 3.67, 95% CI: 0.60 ∼ 22.30; exact P = 0.22), whereas males showed no directional change (paired OR = 1.00, 95% CI: 0.10 ∼ 9.61; exact P = 1.00). The female-to-male ROR was 3.67 (95% CI: 0.20 ∼ 66.31), with wide confidence intervals reflecting limited discordant transitions.

For the bias, the LMM revealed a significant three-way interaction with a medium effect size among sex, group, and timepoint (F1,22 = 4.95, p = 0.037, η²p = 0.18, ω²p = 0.14). The model also identified a significant main effect of sex, with a medium effect size (F1,22 = 4.87, p = 0.038, η²p = 0.18, ω²p = 0.14). Decomposition via simple two-way interaction analyses showed a significant interaction between group and timepoint with a strong effect size for female rats (F1,10 = 9.20, p = 0.013, η²p = 0.48, ω²p = 0.41), but not in male rats (F1,12 = 0.00, p = 1.000, η²p = 0.00, ω²p = 0.00; *Figure 4A*). Subsequent simple main effects of timepoint within each sex and group stratum were not significant (*Figure 4B*).

**Figure 4.**
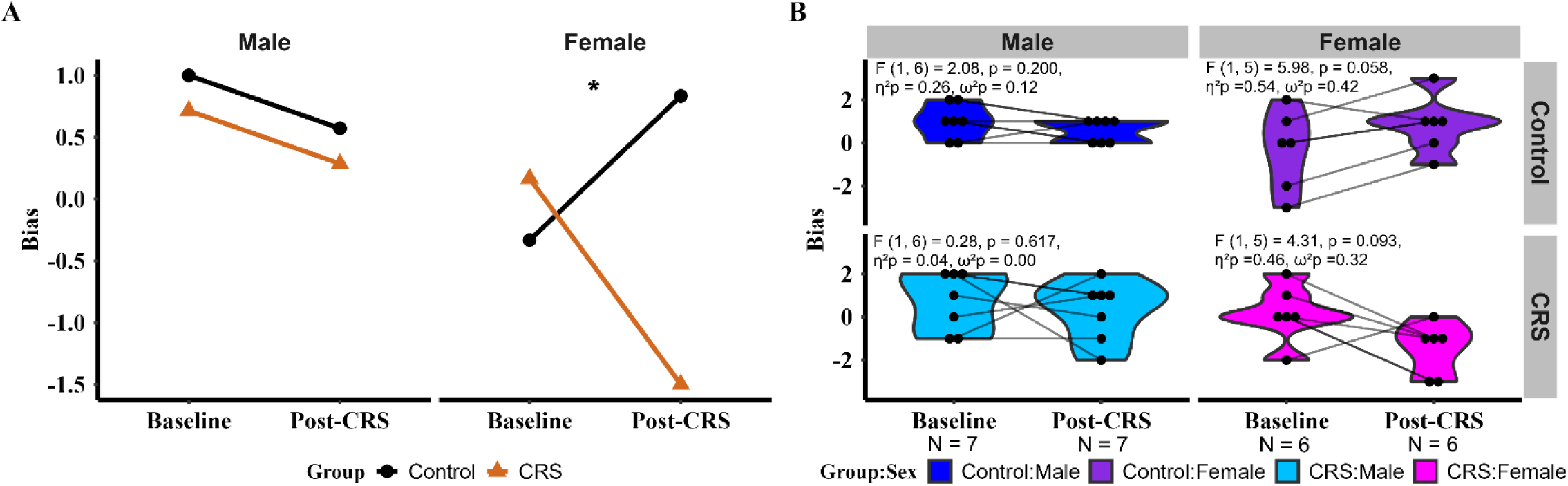
Plots Showing Findings from LMM for Bias. **A).** Interaction plot showing the interaction between group and timepoint was significant for female rats, but not for male rats; **B).** Violin plots showing no significant differences in the bias at baseline and post-CRS, for each sex-by-group stratum. Each paired set of observations is connected by a line.

### Other Behavioral Measures

For response accuracy, probe duration and OFT measures, LMM models did not identify significant three-way interactions among sex, group, and time (*Supplementary Spreadsheet*). However, significant main effects of sex were identified for the probe duration and OFT measures, but not for response accuracy. For instance, there was no significant three-way interaction (F1,22 = 0.94, p = 0.34, η²p = 0.04, ω²p = 0.00), but a significant main effect of sex (F1,22 = 6.47, p = 0.02, η²p = 0.23, ω²p = 0.19) for probe duration.

### Network Connectivity

NBR analysis identified a nine-edge subnetwork associated with the sex × group × timepoint interaction (*Figure 5A)*. These nine edges spanned subcortical, cerebellar, brainstem, and cortical networks, with multiple edges involving the ventral striatum–cortical and spatial attention networks (Table 1). This subnetwork was significant by size (pFWE = 0.04; *Figure 5B*), but not by strength (pFWE = 0.11; *Figure 5C*). Further decomposition revealed five edges within the subnetwork that were associated with a significant interaction between group and timepoint in female rats (pFWE = 0.02). There five edges (edge 21, 26, 35, 50 and 76) were interconnected through two primary networks: auditory thalamus–brainstem and lateral–temporal cortical networks (Table 1). Subsequent simple main effects of timepoint within each sex and group stratum were not significant for the subnetwork.

**Figure 5.**
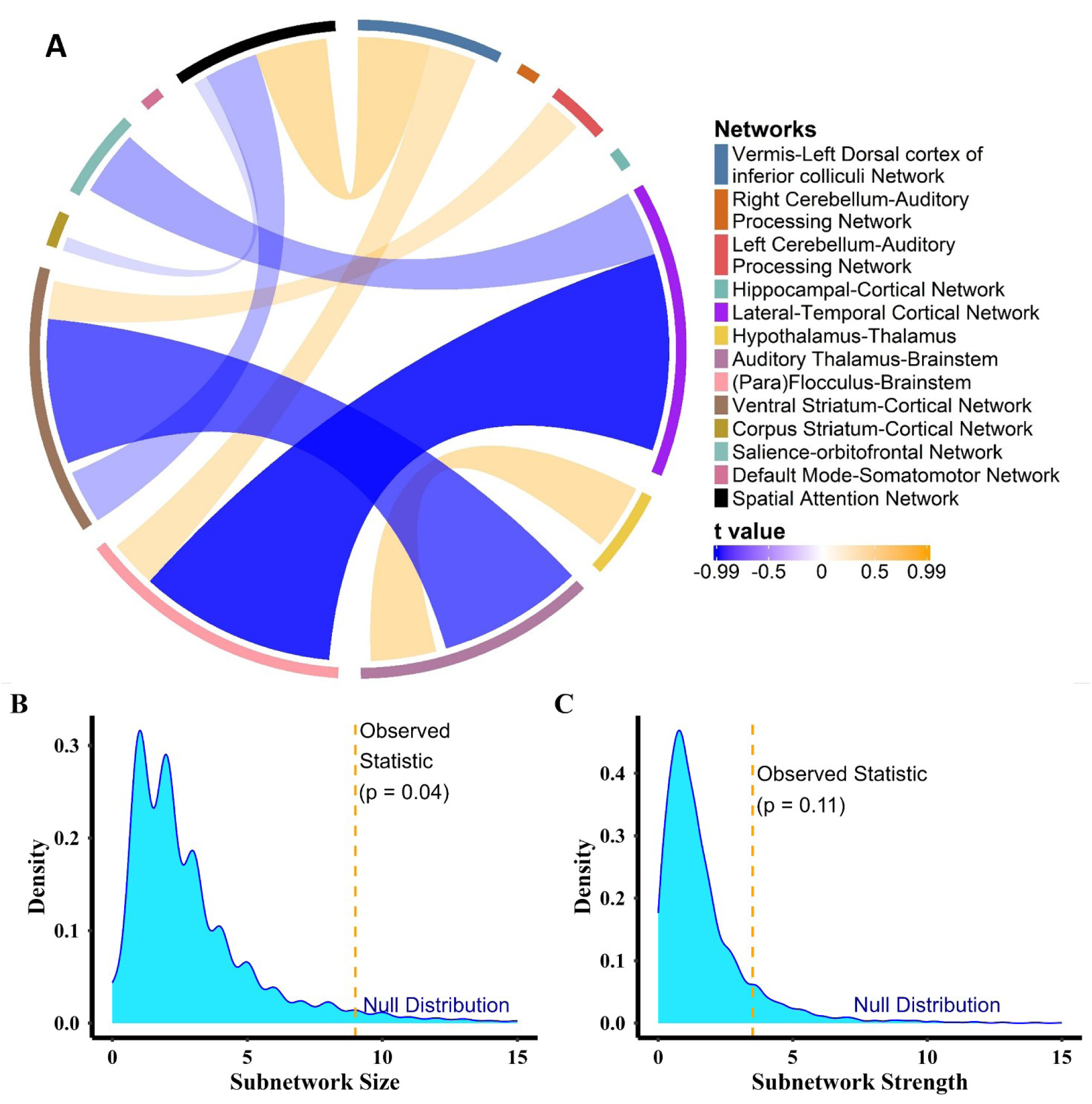
Plots Showing Findings from NBR Analysis. **A).** Chord diagram showing the nine-edge subnetwork identified by NBR analysis. Ribbon color indicates the sign and magnitude of the edge-level test statistic; **B).** Density plot showing null distribution of maximal subnetwork size over 5000 permutations. The dashed line indicates the significant size, corrected for FWE (p = 0.04)**; C).** Density plot showing null distribution of maximal subnetwork strength over 5000 permutations. The dashed line indicates the observed subnetwork strength, which did not survive FWE corrections (p = 0.11).

**Table 1.** Edges Comprising the Subnetwork Identified by the NBR Analysis.

| ID* | Edges |  | t values |
| --- | --- | --- | --- |
|  | Network 1 | Network 2 |  |
| 21 | *Hypothalamus–Thalamus* | *Auditory Thalamus–Brainstem* | 0.33 |
| 22 | Vermis–Left Dorsal cortex of inferior colliculi Network | (Para)Flocculus–Brainstem | 0.24 |
| 26 | *Lateral–Temporal Cortical Network* | *(Para)Flocculus–Brainstem* | -0.99 |
| 31 | Left Cerebellum–Auditory Processing Network | Ventral Striatum–Cortical Network | 0.19 |
| 35 | *Auditory Thalamus–Brainstem* | *Ventral Striatum–Cortical Network* | -0.73 |
| 50 | *Lateral–Temporal Cortical Network* | *Salience–orbitofrontal Network* | -0.33 |
| 67 | Vermis–Left Dorsal cortex of inferior colliculi Network | Spatial Attention Network | 0.36 |
| 75 | Ventral Striatum–Cortical Network | Spatial Attention Network | -0.27 |
| 76 | *Corpus Striatum–Cortical Network* | *Spatial Attention Network* | -0.07 |

*Note: \* Edge ID indicates the original ordering of each edge in the upper triangle of the full connectivity matrix (13 × 13). Details of these networks can be found in Dai et al. (28).*

## Discussion

The present study aimed to determine whether chronic stress during adolescence was associated with sex-dependent alterations in cognitive bias and functional connectome in emerging adulthood. A major innovation of the work lies in the longitudinal integration of a translationally relevant judgment bias assay with resting-state connectomes, which allows behavioral and network trajectories to be examined in parallel within the same cohort. The findings indicate that the change in bias and resting-state reorganization from baseline to post-CRS were sex-dependent. Specifically, females showed a greater tendency toward negative bias than males, and network analysis identified a subnetwork spanning brainstem, limbic, striatal, insular, parahippocampal, orbitofrontal, and retrosplenial regions. Together, these results highlight the translational importance of considering sex and distributed network organization when modelling how adolescent stress may shape later vulnerability to depression.

### Negative Bias: Longitudinal Effects of Adolescent CRS in Adult Females and Males

The primary behavioral finding of this study was that the directional change in ambiguity appraisal from baseline to post-CRS differed between females and males, as indicated by a significant three-way interaction and a subsequent two-way interaction in females. However, the absence of significant simple main effects within the female CRS group calls for a cautious interpretation. These data do not suggest a uniform transition toward negative bias in females after chronic stress. Rather, the significant interactions indicate that the female CRS and control groups differed in how ambiguity appraisal changed from baseline to post-CRS, whereas no comparable pattern was evident in males. This interpretation is consistent with the paired transition analysis suggesting a greater tendency toward negative bias in females than in males. Importantly, because neither task performance nor anxiety-like behavior showed significant interactions or simple main effects of time, this tendency toward negative bias is less likely to reflect a broad behavioral disruption and more likely to reflect a selective change in ambiguity appraisal. This result is also unlikely to be explained by normative maturation alone, given that both control and CRS groups were assessed in early adolescence and again in young adulthood.

Broadly, the present findings are consistent with evidence that sex differences in cognitive vulnerability to depression become more apparent during adolescence (10). One plausible mechanism is the sex-specific programming of the HPA axis during this period (18). Specifically, repeated glucocorticoid signaling in females may alter stress reactivity, subsequently biasing the appraisal of ambiguous stimuli toward more negative interpretations. In this respect, the present study extends work in adult models (43, 44). Although stress exposure in adulthood can acutely induce negative bias (45), these effects are often transient (46). This raises the possibility that stress during adulthood may be insufficient to recalibrate cognitive appraisal, whereas adolescence represents a critical period for the programming of adult affective processing (18, 47). The current work also extends the scope of the latent vulnerability framework (48), aligning with convergent human and developmental evidence suggesting that a negative interpretation bias under uncertainty is a sex-dependent recalibration of affective systems, gated by the intersection of sex and developmental timing (49, 50).

### Network Connectivity: Longitudinal Effects of Adolescent CRS in Adult Females and Males

The network analysis showed an interaction pattern similar to that observed for bias, and follow-up decomposition localized the female-specific interaction to a five-edge subnetwork. Because this subnetwork was significant by size rather than strength, the effects of adolescent CRS appear to have differed between females and males at the level of widespread network reorganization rather than connectivity magnitude. This finding is consistent with prior evidence showing that stress can reorganize large-scale brain networks (51, 52) and broadly aligns with current models of depression that increasingly emphasize abnormalities across interacting large-scale networks rather than dysfunction within a single affective circuit (53, 54). The two primary networks highlighted in the follow–up analysis point to resting-state networks associated with sensory regulation (55), multisensory integration (56) and contextual association (57, 58). Within the identified subnetwork, the thalamus likely mediates sensory gating (55), whereas the para–hippocampal regions, including the perirhinal, postrhinal, and entorhinal cortices, support contextual and associative processing (56). Furthermore, the insula and sensory association cortices may provide the necessary architecture for multisensory and interoceptive integration (57, 58). Beyond these two primary networks, the subnetwork also extends to functional networks associated with physiological state (59), motivational salience (60–62), and spatial attention (63). Specifically, the hypothalamus–thalamus network likely regulates physiological state through reciprocal connections. The hypothalamus monitors internal and external conditions to influence autonomic, endocrine, and behavioral responses, whereas the thalamus contributes to sensory and emotional processing. The ventral striatum–cortical, corpus striatum–cortical, and salience–orbitofrontal networks are likely engaged in motivational salience. The corticostriatal circuits serve as a high-level integrative system that links cortical sensory, emotional, and cognitive input to behavioral prioritization and action selection (60). The salience network and orbitofrontal cortex form a coordinated system for the prioritization and appraisal of environmental cues (61, 62). The dorsal hippocampus and retrosplenial cortex constitute the core functional components of the spatial attention network (28), which likely supports the selective allocation of attention across space, thereby guiding adaptive behaviors (63).

This functional profile is relevant to stress because mounting evidence suggests that chronic stress can alter brain organization beyond classical affective circuits, such as the prefrontal-limbic circuit. For example, a longitudinal imaging study shows structural and functional changes across the thalamus, hypothalamus, striatum, brainstem, auditory cortex, insula, and entorhinal cortex as stress exposure unfolds over time (51). Importantly, sex differences in neural response to adolescent stress have been reported, with the clearest evidence pointing to sex-dependent alterations in the hippocampus (64, 65), amygdala (66, 67), and hypothalamus (68, 69). More limited but growing evidence also supports the involvement of insula-prelimbic pathway (70), ventral striatum (71) and brainstem dopaminergic nuclei such as the ventral tegmental area (72). However, direct evidence for sex-dependent effects of adolescent stress remains limited for other regions within the subnetwork identified here, including the thalamus, auditory cortex, and entorhinal cortex. Further animal and human studies are therefore needed to determine whether these regions also contribute to the sex differences in neural response to adolescent stress.

### Overall Sex Differences in Bias, Training and Probe Duration

The significant main effect of sex on bias and probe duration, together with longer training duration in female Wistar rats, is also reported in Sprague-Dawley rats (46, 73, 74). The absence of sex differences in response accuracy makes it less likely that the overall sex differences in bias, training and probe duration reflect poorer task learning or maintenance in one sex. Instead, these findings suggest a broader sex difference in response strategies associated with choice, learning, and response speed (75, 76).

### Limitations and Future Direction

Several limitations of this study should be acknowledged.

First, given the modest sample size and the absence of a prior power analysis, the findings on bias and resting-state networks require cautious interpretation. Although both measures showed significant sex-dependent interactions, simple main effects of timepoint within each sex-by-group stratum were not significant. This may reflect limited statistical power after stratification, particularly for the MRI analyses. Longitudinal studies with larger sample sizes are warranted to determine whether these sex-dependent effects of adolescent CRS represent reliable changes in behavior and network organization.

Second, the study applied one CRS protocol without physiological measures of stress response. Equivalent restraint schedules do not guarantee equivalent biological stress load across sexes (16). In addition, adolescent stressors vary in intensity, predictability, and social context, with potentially different behavioral and neural outcomes in adulthood. Future work should examine whether the present findings generalize across different adolescent stressors, while incorporating physiological indices of stress response, such as corticosterone and adrenal measures.

Third, the anesthetized “resting” state during rs-fMRI captures stable network reorganization, whereas the awake “deciding” brain reflects dynamic cognitive processing (77, 78). Task performance is likely to depend on multiple processes, including sensory discrimination, selective attention, memory for learned cue–reward contingencies, and the transformation of ambiguous sensory information into action (79). Future investigations should employ multiphoton imaging or electrophysiology to record real-time neural activity during the task (80, 81), directly assessing how circuit dynamics give rise to individual choices under ambiguity.

## Conclusion

This study suggests that adolescent chronic restraint stress in rodents is associated with sex-dependent alterations in cognitive bias and distributed resting-state networks. The findings highlight the importance of sex and network organization in understanding how adolescent stress may shape later vulnerability to depression.

## Supporting information

Supplementary Scripts

## Acknowledgments

The authors thank Lakshini Piyasiti, Neha Nair, Rebecca Goodyear, and Will Copping for assistance with chronic restraint stress, and UWA Animal Care Services for excellent animal care. The authors also acknowledge the facilities and scientific and technical assistance of the National Imaging Facility, a National Collaborative Research Infrastructure Strategy (NCRIS) capability, at the Centre for Microscopy, Characterisation and Analysis, UWA.

## CRediT authorship contribution statement

**Twain Dai:** Writing - Original Draft, Writing - Review & Editing, Conceptualization, Methodology, Software, Formal analysis, Investigation, Resources, Data Curation, Visualization, Project administration; **Liz Jaeschke-Angi:** Investigation (MRI acquisition)**, Marissa Penrose-Menz:** Investigation (MRI acquisition)**, Tim Rosenow:** Investigation (MRI acquisition)**, Jennifer Rodger:** Conceptualization, Writing - Review & Editing, Supervision.

