## Supplementary Table for "Sex-Dependent Effects of Chronic Stress During Adolescence on Cognitive Bias and Functional Connectome in Young Adult Rats"

**SUPPLEMENTARY INFORMATION**

**Sex-Dependent Effects of Chronic Stress**

**During Adolescence on Cognitive Bias and**

**Functional Connectome in Young Adult Rats**

Dai *et al.*

Methods

**Supplementary Table 2. Animal MRI parameters for structural MRI and rs-fMRI acquisition**

| **Parameters** | **T2w sMRI** | **rs-fMRI (EPI)** |
| --- | --- | --- |
| **Repetition Time** (ms) | 1000 | 1000 |
| **Echo Time** (ms) | 6.7 | 15 |
| **Spatial Resolution** | 0.15 × 0.15 × 0.3 mm^3^ | 0.3 × 0.3 mm^2^ |
| **Field of View** | 28.8 × 15 × 27.3 mm^3^ | 27.0 × 17.4 mm^2^ |
| **Slice Thickness** (mm) | 27.3 | 1.0 |
| **Volume** | 1 | 1000 |
| **Effective Acquisition Bandwidth** (Hz) | 96153.8 | 250000 |
| **Flip Angle** (Degree) | 90 | 53 |
| **Fat Suppression** | Yes | Yes |
| **Read Orientation** | Left to Right | Left to Right |
| **Scan Time** | 22 min 40 s | 16 min 40 s |

**Supplementary Table 1. Judgement Bias Task Combinations**

| **Task Combination** | **High Rewards (Three Cheerios)** | | |  | **Low Rewards (Half Cheerio)** | | |
| --- | --- | --- | --- | --- | --- | --- | --- |
|  | **Location 1** | **Scent 1** | **Tactile 1** |  | **Location 2** | **Scent 2** | **Tactile 2** |
| **Combination 1** | Left | Cinnamon | Coarse |  | Right | Coriander | Fine |
| **Combination 2** | Left | Coriander | Coarse |  | Right | Cinnamon | Fine |
| **Combination 3** | Left | Cinnamon | Fine |  | Right | Coriander | Coarse |
| **Combination 4** | Left | Coriander | Fine |  | Right | Cinnamon | Coarse |
| **Combination 5** | Right | Cinnamon | Coarse |  | Left | Coriander | Fine |
| **Combination 6** | Right | Coriander | Coarse |  | Left | Cinnamon | Fine |
| **Combination 7** | Right | Cinnamon | Fine |  | Left | Coriander | Coarse |
| **Combination 8** | Right | Coriander | Fine |  | Left | Cinnamon | Coarse |
