## Supplementary Scripts for "Sex-Dependent Effects of Chronic Stress During Adolescence on Cognitive Bias and Functional Connectome in Young Adult Rats"

```
###The script is written by Twain Dai.
library(dabestr)
library(tidyverse)
library(dplyr)
library(tidyr)
library(openxlsx)
library(str2str)
library(R.utils)
library(data.table)
library(parallel)
library(glmnet)
library(foreach)
require(vip)
library(caret)
library(ggalluvial)
library(nestedcv)
library(Boruta)
library(pROC)
library(effectsize)
library(rstatix)
library(lme4)
library(lmerTest)
library(performance)
library(ez)
library(afex)
library(BayesFactor)
library(ggpubr)
library(rmcorr)
library(stringr)
library(sf)
library(sp)
library(imputeTS)
library(ggmap)
library(cowplot)
library(corrplot)
library(keras3)
library(tensorflow)

setwd("C:/Users/Twain/CAB_2024/Animal_OFT_videos")
source('C:/Users/Twain/CAB_2024/Animal_OFT_videos/DLCAnalyzer_Functions_final.R')
source("nbr_lme_subset.R")

##OFT automation

#Remove irrelevant columns and save as new DLC files
input_folder <- "C:/Users/Twain/CAB_2024/Animal_OFT_videos/DLC_Outputs"
output_folder <- "C:/Users/Twain/CAB_2024/Animal_OFT_videos/OFT_CSV/"
files <- list.files(input_folder,pattern =
"*DLC_resnet50_CAB2024Jul27shuffle1_4000000\\.csv$")
for (i in files){
  OFT_csv <- read.csv(paste(input_folder,i,sep = "/"))
  OFT_csv <- OFT_csv %>% select(-c(38:49))
  colnames(OFT_csv) <- sub("\\..*", "",colnames(OFT_csv))
  filename <- gsub("DLC_.*$","",i)
  filename <- gsub(" ", "", paste(filename,".csv"))
  filepath <- file.path(output_folder, filename)
  write.csv(OFT_csv, file = filepath, row.names = FALSE)
}

input_folder <- "C:/Users/Twain/CAB_2024/Animal_OFT_videos/OFT_CSV/"
files <- list.files(input_folder)

pipeline <- function(path){
  Tracking <- ReadDLCDataFromCSV(path, fps = 30)
  Tracking <- CalibrateTrackingData(Tracking, "area",in.metric = 50*50, points = c(
  "Topright","Topleft","Bottomleft","Bottomright"))
  Tracking <- CleanTrackingData(Tracking, likelihoodcutoff = 0.95)
  Tracking <- CutTrackingData(Tracking, keep.frames = c(0:17999))
  Tracking$median.data <- Tracking$median.data %>%
                          add_row(PointName = c("Centertopleft","Centertopright",
                          "Centerbottomleft", "Centerbottomright"),
                                  x = c(((Tracking$median.data$x[9]-Tracking$
                                  median.data$x[8])/5 + Tracking$median.data$x[8]),
```

```
                                                                  (4*(Tracking$median.data$x[9]-Tracking$median.data$x[8])/5 + Tracking$median.data$x[8]),
                                                                  ((Tracking$median.data$x[10]-Tracking$median.data$x[11])/5 + Tracking$median.data$x[11]),
                                                                  (4*(Tracking$median.data$x[10]-Tracking$median.data$x[11])/5 + Tracking$median.data$x[11])),
                                                            y = c(((Tracking$median.data$y[11]-Tracking$median.data$y[8])/5 + Tracking$median.data$y[8]),
                                                                   ((Tracking$median.data$y[10]-Tracking$median.data$y[9])/5 + Tracking$median.data$y[9]),
                                                                   (4*(Tracking$median.data$y[11]-Tracking$median.data$y[8])/5 + Tracking$median.data$y[8]),
                                                                   (4*(Tracking$median.data$y[10]-Tracking$median.data$y[9])/5 + Tracking$median.data$y[9])))
  row.names(Tracking$median.data) <- Tracking$median.data$PointName
  Tracking <- AddZones(Tracking,z = data.frame(center = c("Centertopleft", "Centertopright", "Centerbottomright", "Centerbottomleft","Centertopleft"),rena = c("Topleft","Topright","Bottomright","Bottomleft","Topleft")))
  Tracking <- AddBinData(Tracking,unit = "minute", binlength = 5)
  Tracking <- OFTAnalysis(Tracking, movement_cutoff = 5, integration_period = 5, points = "Bodycenter")
  Tracking$Report<- c(Tracking$Report,ZoneReport(Tracking, point = "Bodycenter", zones = c("center"), zone.name = "Bodycenter.periphery", invert = TRUE))
  return(Tracking)
}
TrackingAll <- RunPipeline(files,input_folder,FUN = pipeline)
Total_Report <- MultiFileReport(TrackingAll)
Total_Report <- Total_Report %>%
  add_column(`Time Bins` = "Total Ten Minutes", .before = 2) %>%
  dplyr::rename("Animal_Timepoint" = "file") %>%
  mutate(Animal_Timepoint = gsub(".csv", "", Animal_Timepoint))

save(TrackingAll, file = "TrackingAll.rda")

cl<- parallel::makeCluster(4)
doParallel::registerDoParallel(cl)
Rat_OFT <- foreach(i = files, .packages = c("sf","sp"),
                          .final = function(i) setNames(i, files)) %:%
  foreach(f = seq_along(TrackingAll[[i]]$frame)) %dopar% {
    x <- c(TrackingAll[[i]]$data[["Neck"]]$x[f],TrackingAll[[i]]$data[["Lefthip"]]$x[f],TrackingAll[[i]]$data[["Righthip"]]$x[f],TrackingAll[[i]]$data[["Neck"]]$x[f])
    y <- c(TrackingAll[[i]]$data[["Neck"]]$y[f],TrackingAll[[i]]$data[["Lefthip"]]$y[f],TrackingAll[[i]]$data[["Righthip"]]$y[f],TrackingAll[[i]]$data[["Neck"]]$y[f])
    Coords <- cbind(x,y)
    st_polygon(list(Coords))
  }


# Extract inner and outer polygon for each video

Center_list <- foreach (i = files, .packages = c("sf","sp"),
                             .final = function(i) setNames(i, files)) %dopar% {
                               Center <- as.matrix(TrackingAll[[i]]$zones$center)
                               st_polygon(list(Center))
                             }

Periphery_list <- foreach (i = files, .packages = c("sf","sp"),
                                .final = function(i) setNames(i, files)) %dopar% {
                                  Center <- as.matrix(TrackingAll[[i]]$zones$center)
                                  Arena <- as.matrix(TrackingAll[[i]]$zones$arena)
                                  Center_sf <- st_polygon(list(Center))
                                  Arena_sf <- st_polygon(list(Arena))
                                  st_difference(Arena_sf, Center_sf)
                                }

Zone <- c("Center", "Periphery")
# Inspect and save animal location in each zone, binary logic (TRUE/FALSE) search.
```

```
Zone_counts <- list()
for (z in Zone){
  Z <- eval(as.name(paste0(z,"_list")))
  Zone_count <- foreach(i = files, .packages = c("sf","sp"),
                        .multicombine = TRUE,
                        .final = function(i) setNames(i, files)) %:%
    foreach(f = seq_along(1:length(Rat_OFT[[i]])),
            .combine = cbind) %dopar% {
              st_within(Rat_OFT[[i]][[f]], Z[[i]], sparse = FALSE)}
  Zone_counts[[z]] <- cbind(files, plyr::rbind.fill(
    lapply(Zone_count,function(y){
      as.data.frame(y,stringsAsFactors=FALSE)
    }))) %>%
    unnest(where(is.list)) %>%
    as.data.frame() %>%
    add_column(Zone = z) %>%
    relocate(Zone, .after = "files") }

parallel::stopCluster(cl)

OFT_counts <- as.data.frame(do.call(rbind,Zone_counts)) %>%
  rename_at(1, ~ "Animal") %>%
  `rownames<-`( NULL )

OFT_counts[is.na(OFT_counts)] <- FALSE

sapply(3:ncol(OFT_counts), function(i) {
  OFT_counts[, i] <<- as.numeric(OFT_counts[, i])
})

T10min <- OFT_counts %>%
  select(1:18002) %>%
  mutate(Animal = gsub(".csv", "", Animal)) %>%
  mutate(Frame = rowSums(across(where(is.numeric)))) %>%
  relocate(Frame, .after= Zone) %>%
  select(1:3) %>%
  spread(key = Zone, value = Frame) %>%
  mutate(T10_Center = Center/30,
         T10_Periphery = Periphery/30) %>%
  select(-2:-3)

F5mins <- OFT_counts %>%
  select(1:9002) %>%
  mutate(Animal = gsub(".csv", "", Animal)) %>%
  mutate(Frame = rowSums(across(where(is.numeric)))) %>%
  relocate(Frame, .after= Zone) %>%
  select(1:3) %>%
  spread(key = Zone, value = Frame) %>%
  mutate(F5_Center = Center/30,
         F5_Periphery = Periphery/30) %>%
  select(-2:-3)

S5mins <- OFT_counts %>%
  select(1:2,9003:18002) %>%
  mutate(Animal = gsub(".csv", "", Animal)) %>%
  mutate(Frame = rowSums(across(where(is.numeric)))) %>%
  relocate(Frame, .after= Zone) %>%
  select(1:3) %>%
  spread(key = Zone, value = Frame) %>%
  mutate(S5_Center = Center/30,
         S5_Periphery = Periphery/30) %>%
  select(-2:-3)

df_list <- list(T10min,F5mins,S5mins)
OFT <- df_list %>%
  reduce(left_join, c("Animal")) %>%
  select(-contains("Center")) %>%
  mutate(T10_Periphery_Percent = T10_Periphery/600,
         F5_Periphery_Percent = F5_Periphery/300,
         S5_Periphery_Percent = S5_Periphery/300) %>%
  separate(Animal, c("ID", "Timepoint", "Date"), "_") %>%
  select(-Date)  %>%
  mutate(Timepoint = case_when(
```

```
    Timepoint %in% "OFT1" ~ "Baseline",
    Timepoint %in% "OFT3" ~ "CRS"))

write.xlsx(OFT, file = "OFT_CAB2024.xlsx",
           col.Names = FALSE, row.Names = FALSE, append = TRUE)


##Statistic Analysis
CAB2024 <- read.xlsx("CAB2024.xlsx", sheet = "CAB_bias")
CAB2024 <- CAB2024 %>%
  mutate(CAB_Pos2Neg = (Large-Small)/2) %>%
  relocate(CAB_Pos2Neg, .after = Bias) %>%
  mutate(Large_accuracy = 1 - Large_wrong_trials/12) %>%
  mutate(Small_accuracy = 1 - Small_wrong_trials/12) %>%
  relocate(c(Large_accuracy, Small_accuracy), .before = Accuracy) %>%
  mutate_at(vars(Large_accuracy, Small_accuracy, Accuracy), round, 4) %>%
  select(-contains("trials"))  %>%
  rename_at('Animal_ID', ~'ID') %>% select(-Probe)

CAB2024 <- CAB2024 %>% filter(Timepoint %in% c("Baseline2","Post-CRS")) %>%
  mutate(Timepoint = case_when(
    Timepoint %in% "Baseline2" ~ "Baseline",
    Timepoint %in% "Post-CRS" ~ "CRS"))

OFT <- read.xlsx("OFT52_CAB2024.xlsx",sep.names = " ") %>% select(-3:-5)

df_list <- list(CAB2024, OFT)
CAB2024_OFT <- df_list %>%
  reduce(left_join, c("ID","Timepoint")) %>%
  mutate_at("CAB_Pos2Neg", as.integer)  %>%
  mutate_at(c("ID","Sex", "Bias", "Timepoint","Group"), as.factor)  %>%
  mutate(Group = factor(Group,levels = c("Control","CRS")))  %>%
  mutate(Sex = factor(Sex,levels = c("Male","Female")))  %>%
  as.data.frame()
contrasts(CAB2024_OFT$Sex) <- matrix(c(-0.5, 0.5), ncol = 1)
contrasts(CAB2024_OFT$Timepoint) <- matrix(c(-0.5, 0.5), ncol = 1)
contrasts(CAB2024_OFT$Group) <- matrix(c(-0.5, 0.5), ncol = 1)


NetworkFC <- read.csv("All_timeseries_13networks.csv")
NetworkFC_COLNAMES <- data.frame(colnames(NetworkFC))
colnames(NetworkFC_COLNAMES)[1] <- "psudo"
NetworkFC_COLNAMES <- NetworkFC_COLNAMES %>% mutate(psudo = str_replace(psudo,
"Network","Network.")) %>%
  separate(psudo, c("Network", "V1", "V2")) %>%
  unite("Edge", Network:V2, sep = "_") %>%
  mutate(Edge = str_replace(Edge, "_NA","")) %>%
  mutate(Edge = str_replace(Edge, "_NA","")) %>% transpose()
NetworkFC_COLNAMES <- as.data.frame(do.call(cbind,NetworkFC_COLNAMES))
rownames(NetworkFC_COLNAMES) <- NULL
colnames(NetworkFC) <- unlist(NetworkFC_COLNAMES)
NetworkFC <- NetworkFC %>%
  separate(ID, c("ID", "Timepoint"))


df_list <- list(CAB2024_OFT, NetworkFC)
CAB2024_OFT_NetworkFC <- df_list %>%
  reduce(left_join, c("ID","Timepoint")) %>%
  mutate_at(c("ID","Timepoint"), as.factor) %>%
  filter_at(vars(Network_1_2), all_vars(!is.na(.))) %>%
  group_by(ID) %>% filter(n() == 2)

cl<- parallel::makeCluster(3)
doParallel::registerDoParallel(cl)

normality <- foreach(i = colnames(CAB2024_OFT[c(5,9:ncol(CAB2024_OFT))]), .combine =
rbind,
                     .packages=c('tidyverse','afex','performance')) %dopar% {
                       tmp <- lmer(eval(as.name(i)) ~ Sex*Group*Timepoint + (1 | ID),
                       CAB2024_OFT)
                       check_normality(tmp) %>%
                         data.frame() %>%
                         add_column(Variable = i)}
```

```r
heteroscedasticity <- foreach(i = colnames(CAB2024_OFT[c(5,9:ncol(CAB2024_OFT))]),.combine = rbind,
                              .packages=c('tidyverse','afex','performance')) %do% {
                                tmp <- aov_ez(id = "ID", dv = i, between = c("Group", "Group"),
                                       within = "Timepoint", observed = "Group",
                                       data = CAB2024_OFT)
                                print(check_heteroscedasticity(tmp)) %>%
                                  data.frame() %>%
                                  add_column(Variable = i)}

LMM_Behav <- foreach(i = colnames(CAB2024_OFT[c(5:7,9:ncol(CAB2024_OFT))]), .combine = rbind,
                     .packages=c('tidyverse','effectsize','lmerTest','rstatix')) %dopar% {
                       tmp <- lmer(eval(as.name(i)) ~ Sex*Group*Timepoint + (1 | ID), CAB2024_OFT)
                       tmp <- anova(tmp, type = 3)
                       tmp %>% parameters::parameters(es_type = c("omega", "eta")) %>%
                         data.frame() %>%
                         mutate(effectsize = interpret_omega_squared(Omega2_partial, rules = "cohen1992")) %>%
                         add_column(Variable = i)}
write.xlsx(LMM_Behav, file = "LMM_Behav.xlsx", rowNames = FALSE, colNames = TRUE)

set.seed(44444)
nbr_result <- nbr_lme(net = CAB2024_OFT_NetworkFC[,-(1:15)], nnodes = 13,
                      idata = CAB2024_OFT_NetworkFC[,1:4], mod = "~ Timepoint*Group*Sex",
                      rdm = "~ 1|ID", nperm = 5000, cores = detectCores() - 8,
                      na.action = na.exclude, nudist = TRUE
)
sel_edges <- c(nbr_result$components$`Timepoint1:Group1:Sex1`[,1])

set.seed(44444)
nbr_bySex <-  foreach(i = unique(CAB2024_OFT_NetworkFC$Sex),
                      .packages=c("NBR", "dplyr", "nlme")) %do% {
                        net <- CAB2024_OFT_NetworkFC %>% filter(Sex == i)
                        nbr_lme_subset(net = net[,-(1:15)], nnodes = 13,
                                idata = net[,c(1,3,4)], edge_idx = sel_edges,
                                mod = "~ Timepoint*Group",
                                cores = detectCores() - 8,
                                rdm = "~ 1|ID", nperm = 5000,
                                component_mode = "fixed_set",
                                na.action = na.exclude)}
names(nbr_bySex) <- unique(CAB2024_OFT_NetworkFC$Sex)

set.seed(44444)
nbr_TimepointFemale <-  foreach(i = unique(CAB2024_OFT_NetworkFC$Group),
                      .packages=c("NBR", "dplyr", "nlme")) %do% {
                        net <- CAB2024_OFT_NetworkFC %>% filter(Sex == "Female") %>%
                          filter(Group == i)
                        nbr_lme_subset(net = net[,-(1:15)], nnodes = 13,
                                       idata = net[,c(1,3,4)], edge_idx = sel_edges,
                                       mod = "~ Timepoint",
                                       cores = detectCores() - 8,
                                       rdm = "~ 1|ID", nperm = 5000,
                                       component_mode = "fixed_set",
                                       na.action = na.exclude)}
names(nbr_TimepointFemale) <- unique(CAB2024_OFT_NetworkFC$Group)

set.seed(44444)
nbr_GroupFemale <-  foreach(i = unique(CAB2024_OFT_NetworkFC$Timepoint),
                            .packages=c("NBR", "dplyr", "nlme")) %do% {
                              net <- CAB2024_OFT_NetworkFC %>% filter(Sex == "Female") %>%
                                filter(Timepoint == i)
                              nbr_lme_subset(net = net[,-(1:15)], nnodes = 13,
                                             idata = net[,c(1,3,4)], edge_idx = sel_edges,
                                             mod = "~ Group",
```

```
                                                  cores = detectCores() - 8,
                                                  rdm = "~ 1|ID", nperm = 1000,
                                                  component_mode = "fixed_set",
                                                  na.action = na.exclude)}
names(nbr_GroupFemale) <- unique(CAB2024_OFT_NetworkFC$Timepoint)

Subnetwork <- as.data.frame(nbr_result$components$`Timepoint1:Group1:Sex1`) %>%
  filter(comp == 1) %>%
  mutate(
    from   = as.integer(`3Drow`),
    to     = as.integer(`3Dcol`),
    tvalue = as.numeric(strn),
    abs_t  = abs(tvalue),
    sign   = ifelse(tvalue > 0, "positive", "negative")
  )
```
